# Ecological and social factors lead to variation in parental care between sexes in a burying beetle

**DOI:** 10.1101/2024.10.22.619607

**Authors:** Donghui Ma, Long Ma, Jan Komdeur

## Abstract

Sexual conflict over parental care represents a divergence in the evolutionary interests between males and females, consequently leading to distinct sex roles in parental care and reproductive strategies. However, whether and how various ecological and social environments influence such sex differences remains largely unclear. In the present study, using the burying beetles *Nicrophorus vespilloides*, we experimentally investigated the impacts of resource availability and inter- and intra-specific competition pressure on individual parental care and reproduction in males and females. We manipulated the resource availability for the breeding pairs by providing them with either large or small mouse carcasses, and constructed the presence of intra- and inter-specific competition, by introducing an additional, small pair of beetles and blowfly maggots *Calliphora*, respectively. We expected that males would decrease their parental care, while females would increase theirs, in response to smaller carcass. We also expected that the presence of intra- and inter-specific competition differentially affected the cares of males and females. We found that, compared to males and females breeding on large carcasses, males did not change their care, whereas females increased their care, when breeding on small carcasses. In the presence of another pair of adults, both males and females increased their parental care, whereas in the presence of blowfly maggots they both decreased their parental care. Our results showed that males and females adjust their parental care based on resource availability and competition pressure, with sex-specific differences driven by ecological and social factors. Our study sheds light on the importance of sex-specific parental care in response to various external drivers which contributes to a better understanding of the evolution of sex conflict over parental care.

## Introduction

Sexual conflict is a fundamental and long-standing topic in behavioural ecology, which refers to an antagonistic selection process that acts on two sexes with different interests (reviewed by Arnqvist & Rowe, 2005). This conflict arises when males and females face unmatched or opposing selection pressures on costly reproductive traits and behaviours (e.g., parental care behaviour), causing variation of optimal fitness benefits between sexes (Parker, 1979; Arnqvist & Rowe, 2005; Bonduriansky *et al*., 2008; Lessells, 2012; Taborsky *et al*., 2014; Paquet & Smiseth, 2016). “Anisogamy” theory has been proposed to explain the different sex roles and their consequent effects on mating and parental care, and it hypothesizes that the asymmetry between sexes may emerge from gamete which may extend to parental investment at the pre- and post-natal stages of offspring (Trivers, 1972). Sexual conflict over parental care can be considered as tug-of-war interactions between sexes, because two parents share benefits from their combined parental effort to their young, whereas the costs of care are paid by each parent due to its own effort to raise the young. Consequently, it could be expected that each parent is under selection to minimise its individual effort by grafting more parental workload over to the other parent (Arnqvist & Rowe, 2005; Houston *et al*., 2005; Székely *et al*., 2014; Paquet & Smiseth, 2016). As a result, two critical questions regarding sexual conflict over parental care have intrigued biologists for decades: 1) whether various forms of parental care evolve within and across animal taxa, ranging from uniparental care to male/female-biased care to biparental care (e.g., Clutton-Brock, 1991; Royle *et al*., 2012; Liker *et al*., 2013; Janicke *et al*., 2016; Vági *et al*., 2019; Gonzalez-Voyer *et al*., 2022), and are facilitated by sexual conflict over parental care, and 2) how sexual conflict over parental care is resolved, which results in biparental care to occur as Evolutionary Stable Strategy (ESS; Harrison *et al*., 2009; Houston *et al*., 2005; reviewed by Paquet & Smiseth, 2016). In this study, we focus on the second question.

Theoretical studies have proposed three potential types of behavioural interactions between male and female parents which may explain the resolution of sexual conflict over parental care (Johnstone & Hinde, 2006; Lessells, 2012; Paquet & Smiseth, 2016), including negotiation, matching and sealed bid responses. Negotiation models assume each parent directly adjusting its level of care in response to its partner’s investment, and that the focal parent may partially compensates for a decreased investment by its partner (McNamara *et al*., 1999). However, matching models highlight a matching strategy deployed by each parent in response to its partner, where each parent adjusts its level of care to match any increase or reduction in the partner’s contribution (Johnstone & Hinde, 2006). Sealed bid models assume that the parental contribution of each individual is fixed and not influenced by its partner (Houston & Davies, 1985). Empirical works across animal species (almost in avian species) provide evidence for all three models, where the two parents deploy similar behavioural strategies to respond to the workload of their partners (Schwagmeyer *et al*., 2002; Johnstone & Hinde, 2006, Harrison *et al*., 2009). For example, females of Great tits *Parus major* fully compensate for their partner’s decrease in feeding rates, while males do not show any compensation and even tend to decrease their feeding rates (Sanz *et al*., 2000). However, whether the alternation of such behavioural strategies is influenced by ecological and social circumstances remains less explored.

Across and within species, evolutionary variation in magnitude and forms of parental care is driven by sex-specific costs and benefits, which is also the result of changes in ecological and social environments experienced by males, females, or both sexes (e.g., van Dijk *et al*., 2012; Zheng *et al*., 2018; Ma *et al*., 2021; Long *et al*., 2022). Normally, individuals adjust their decisions in parental care according to the variation of internal (e.g., body condition, age and the level of hormones; Achorn & Rosenthal, 2020) and external factors (i.e., ecological and social environments; Ma *et al*., 2021). For example, parents with good body conditions are likely to invest more time and energy to care for their offspring, gaining greater reproductive benefits, compared to those with bad body conditions (Achorn & Rosenthal, 2020). This is because bad body conditions may lead to constrained resources available for self-maintenance and future breeding opportunities (Soulsbury, 2019; Pontzer & McGrosky, 2022). As one of external factors, resource availability, such as food abundance and territory quality, affects individual parental behaviour and reproduction. While breeding in territory of good quality, individuals that forage easier and could gain enough food resources for breeding tend to increase their parental care for their offspring, compared to those who breed in a lower quality territory (Gauthier & Jong, 2021). As another external factor, the pressure of competition with inter- and intra-specific individuals is found to directly influence an individual’s reproductive success due to limited resources for reproduction, which also alters an individual’s parental behaviour and investment (e.g., Requena *et al*., 2009; Grayson *et al*., 2013; Chan *et al*., 2019; Ratz *et al*., 2022). Nevertheless, it remains less understood whether males and females differentially respond to different ecological and social environments by adjusting their parental investment, and whether such difference in behavioural adjustment is also adaptive and different for males and females.

In this research, we explore the impacts of ecological and social factors on male and female parental care behaviour, and whether and how each parent show plasticity in behaviour in response to the presence of its breeding partner in the different contexts in the burying beetles *Nicrophorus vespilloides*. Burying beetles typically exhibit intricate parental care, which provides an ideal biological system for studying the mechanisms of the co-evolution of parental care between sexes across animal species, especially in insects, and their roles in response to the changes of environmental and social stressors (Paquet & Smiseth, 2016). Burying beetles use small vertebrate carcasses as breeding and food resources that support food for offspring and parents themselves (Scott, 1998). These beetles often breed as pairs, jointly preparing the carcasses and preserving them with anti-microbial secretion (i.e., indirect care; Scott, 1998; Smiseth *et al*., 2005; Steiger *et al*., 2007). Females lay eggs in the soil near the prepared carcass, and then provide care for the developing larvae along with the males. Generally, there occurs a division of parental care between the sexes: males primarily defend the broods against intra- and interspecific competitors and usually leave the carcass earlier than females, whereas females often provide direct post-hatching care, mainly by feeding the larvae on the prepared carcass (e.g., Smiseth *et al*., 2005; Lee *et al*., 2014; Paquet *et al*., 2017; Ratz *et al*., 2020). Carcasses are ephemeral, unpredictable resources for beetles, as the availability of carcass and its size highly determine reproductive success of beetles and influence their parental investment to current reproduction (e.g., Richardson & Smiseth, 2019; Wang *et al*., 2022). Furthermore, it is well documented that other factors also impact the male and female parental care, such as their body size and nutritional condition (e.g., Pilakouta *et al*., 2015; Richardson & Smiseth, 2019), loss of mate (Smiseth *et al*., 2005), brood size (Wang *et al*., 2021), intraspecific interactions with other breeding pairs (Ma *et al*., 2022) and interspecific interactions with mites (De Gasperin *et al*., 2015).

In burying beetles, we can easily manipulate the levels of resource availability by giving beetles carcasses of different size, and created the absence or presence of intraspecific competition by controlling the number of adult beetles in the breeding attempts, and different interspecific competition by introducing different numbers of blowfly maggots that are the main competitors in natural breeding events (Bartlett, 1987a; Scott, 1998; Chan *et al*., 2019). Here, we constructed three experimental groups and one contrasted group to explore whether and how resource availability, intra- and interspecific competitions differentially affect individual parental care behaviour, and whether alternative parental care strategies occur between sexes in different ecological and social contexts. Females typically provide more direct care but take less time on carcass defence and maintenance compared to males (e.g., Eggert *et al*., 1998; Müller *et al*., 1998; Smiseth *et al*., 2005; Wang *et al*., 2021, 2022). However, when breeding on small carcasses, there is a high level of conflict over resources and parental care between females and males, and males may decrease their parental care. When pairs breed on small carcasses, there is increased competition for food between parents and offspring, as well as among siblings, but a decreased workload of carcass maintenance and defence against intruders. This underpins the evolution of a female uniparental care system in this species (e.g., Scott, 1990; Boncoraglio & Kilner, 2012). Given this, we predicted males that utilize small carcasses for reproduction decrease its parental care, whereas their female partners increase parental care. Furthermore, males and females would kill larvae unless their own first larva hatched in cross-fostered broods (infanticide; Bartlett, 1987b; Müller & Eggert, 1990). This suggests that both males and females would benefit from spending more time accessing to the carcass in a joint brood (i.e., cobreeding or communal breeding; Komdeur *et al*., 2013; Ma *et al*., 2022), which allows them to prevent infanticide from other adults, care for their offspring into ensuring current reproductive success, and feed themselves to pursue future reproductive opportunity. Thus, when intraspecific competition is present during breeding events, we expect that both males and females would increase their level of care to offspring. However, in the presence of interspecific competition, we expect males to decline parental care because they will invest more in resource defence, and as a consequence we expect females to increase their parental care due to reduced male care (Trumbo, 2006, 2022). We expected that male and female parents would deploy sex-dependent parental strategies by adjusting parental care in response to their breeding partners, and such parental strategies are altered in different ecological and social contexts. Our research will contribute toward a better understanding of the evolution of parental care and help to understand the eco-evolutionary factors that facilitate sexual conflict.

## Materials and methods

### Burying beetle husbandry

Adult beetles used for this study descended from F_2_ to F_5_ generations of an outbred laboratory population at the University of Groningen. The laboratory population was derived from wild-caught beetles at the estate De Vosbergen, Eelde, the Netherlands (53°08’ N, 06°35’ E), from May to August in 2017 and 2018, using pitfall traps baited with chicken livers. During the entire rearing period, up to six same-sex adult beetles derived from the same brood were kept in rearing boxes (10 cm L × 6 cm W × 8 cm H) with 2 cm depth of wet peat. They were fed twice a week with 1 - 2 mealworms per beetle. Throughout the experiment, all adults and offspring were maintained at a temperature of 20 ℃ with a 16 h: 8 h light: dark photoperiod.

### Experiment protocol

In this study, we designed three independent experimental groups and a control group, to explore the effects of resource availability, and inter- and intra-specific competition on the degree of sexual conflict over parental care in burying beetles. Sexually mature, virgin adult beetles, aged approximately two weeks old at post-eclosion, were selected for this experiment. Just before the experiment, body size of each individual was recorded by measuring the pronotum width using electric callipers (accuracy: 0.001mm) (Beeler *et al*., 2002; Hopwood *et al*., 2016), and weight measured using electric balance (accuracy: 0.001g). Males and females were individually marked by slightly pricking two small holes in one of the elytra (i.e., the left side for male and the right side for female) using an insect pin. This method allowed us to recognise the parental individuals during the subsequent daily inspections of the breeding boxes. For each breeding event, each beetle was paired with a similarly sized, unrelated partner, and housed with its mate into a rearing box with clean peat for 12h, to make sure that females had been fertilized (Ma *et al*., 2022; Wang *et al*., 2021, 2022). Then, each pair was transferred into a breeding box (23 cm L× 19 cm W× 12.5 cm H) with 3∼5 cm of moist peat, and given a thawed, previously frozen, dead mouse as breeding resource.

To create groups with high and low resources availability, beetle pairs were given a larger carcass (ca. 25 gram) or a smaller carcass (ca. 15 gram). These carcass masses were selected as these carcasses could be used by this species under both natural and laboratory conditions (Scott, 1998; Smiseth & Moore, 2008). The small-carcass group was assigned as our resource-availability treatment (*N* = 47), while the large-carcass group as contrasted treatment for the three experimental groups (*N* = 52). To create intraspecific competition treatment (*N* = 78), we placed an additional pair of beetles as intruders at 12 h after the *residential* pair was provided with a ca. 25 gram carcass for breeding. These additional pairs were smaller in body size (approx. 10 – 15% size difference) compared to the resident breeding pairs. This could potentially decrease the harmful risks from intruders to the residents and the brood, such as the usurpation of carcasses by intruders, because an individual’s body size largely determines its competitive ability and intense fights were less likely to occur between opponents with a larger size difference in burying beetles (Beeler *et al*., 2002; Komdeur *et al*., 2013). To construct the interspecific competition treatment (*N* = 33), we created the ‘blowfly maggots usurped’ carcass (ca. 25 gram), by making a small hole (1cm of diameter) on a thawed mouse carcass, and then introducing 2.0 ± 0.2 g newly-hatched blowfly maggots (*Calliphora*) into the carcass. To maintain a continuous pressure of interspecific competition, introduction of 2.0 ± 0.2 g newly hatched blowfly maggots into carcasses was repeated at 24 hours after the onset of each experiment.

### Behaviour observation and data collection

Across all treatment groups, parental care behaviour and body mass change of each beetle individual and reproductive success were measured. During the entire period of each breeding event (from the onset of experiment until larval dispersal from the carcass), we checked each box three times daily for carcass preparation and parental care (07:00–09:00 am, 14:00–16:00 pm, 21:00–23:00 pm, 5-h intervals) by visual inspection (instant scanning) through carefully removing the surface soil of the carcass (Ma *et al*., 2022; Wang *et al*., 2021, 2022). The presence of each beetle on or in the carcass is an indicator of parental care in burying beetles (Ma *et al*., 2022; Wang *et al*., 2021, 2022). We recorded the amount of parental care (only the larger pair as focal parents in the interspecific competition group; Ma *et al*., 2022) during the parental period. This was done by recording at every visual inspection whether a target parent was present on or inside the carcass (i.e., parental care), or whether it was invisible in the soil (i.e., no parental care) (Wang *et al*., 2021, 2022). We calculated the amount of parental care as the number of times the parent was present on the carcass of the total checked times. Once the larvae dispersed, all alive parents were weighed to calculate the body mass change of adults over the breeding events (body mass change = [final mass – initial mass]/ initial mass), which is seen as an indicator of parental investment towards current reproduction (Wang *et al*., 2021, 2022). We also recorded the number of dispersed larvae (hereafter brood size) and weighed them in broods (hereafter brood mass), and the number of newly eclosed beetles after pupation (hereafter offspring number) for evaluating the reproductive and offspring performance of each brood.

### Data analyses

All statistical tests were performed using R version 4.2.1 (R core team, 2022) loaded with packages *car* (Fox & Weisberg, 2019), *lme4* (Bates *et al*., 2015), *MASS* (Venables & Ripley, 2002), *multcomp* (Hothorn *et al*., 2008)*, emmeans* (Lenth, 2023).We used a *cbind* function to combine times of male (i.e., T _male_) or female (i.e., T _female_) being present on or in the carcass, with the times that we checked the breeding boxes (i.e., T _check_) subtracting times of the parent being present on or in the carcass as response variables (i.e., for male: *cbind* (T _male_, (T _check_ – T _male_)); for female: *cbind* (T _female_, (T _check_ –_female_))). These response variables were then connected with explanatory variables using a generalised linear model (GLM) with a binomial error distribution in which the explanatory variables include our contrasted and experimental treatments (i.e., one contrasted group and three experimental groups), spouse care ((T _male_ /T _check_) or (T _female_ /T _check_)), and their interaction. This is to test whether and how male and female individuals respond to the experimental manipulation and their mates’ investment. We also performed a binomial generalised linear mixed model (GLMM) to compare the care of male and female under our treatments, in which we set the treatments, parental sexes and their interaction as fixed factors, and the breeding box ID as a random factor. A similar GLM model was performed for the total parental care of a pair, where the response variable followed the formula: *cbind* ((T _male_ + T _female_), (2 × T _check_ – T _male_ – T _female_)), and the explanatory variable is our contrasted and experimental treatments. Furthermore, the ratio between male and female parental investment (i.e., male care / female care) was also tested using GLM fitted with binomial distribution, in which this ratio is performed as response variable following the formula: *cbind* (T _male_, (T _male_ + T _female_ - T _male_)), and the explanatory variable is our contrasted and experimental treatments. We used negative binomial GLMs to test brood size and offspring number (integer values), and linear (mixed) models to test brood mass (LMs) and parental body mass change (LMMs). We set the contrasted and experimental treatments, male and female care (i.e., (T _male_ /T _check_) and (T _female_ /T _check_)) as explanatory variables. Furthermore, in the models for parental body mass change, we also included genders of parents, and its interaction with our treatments as explanatory variables, and the breeding box ID as random effect. We used the *emmeans* function to do *post hoc* Tukey correction tests when the interaction effects and the treatments showed a significant effect on targeted response variables. Otherwise, we applied the *summary* function to show and report results of models (including those without an interaction), because we only aimed to test the variation of targeted response variables between the three experimental treatments and the contrasted treatment. Figures were generated using the *ggplot2* package (Wickham, 2016).

## Results

### Parental coordination over care between sexes based on resource availability and intra- and interspecific competition

In general, the result of the GLM analysis showed that parental coordination over care differed between treatment groups (χ² = 245.45, *P* < 0.001). Specifically, compared to the contrasted group, a significantly higher total amount of parental care by pairs was observed in the resource-availability and the intraspecific competition groups, and a lower total amount of parental care by pairs in the interspecific competition group (Table 1, Fig. 1a). From the perspective of both male and female individuals (i.e., results of the GLMM model), they significantly adjusted their care in response to the treatments (χ² = 71.92, *P* < 0.001), with females consistently providing a significantly higher level of care than males (χ² = 202.85, *P* < 0.001; Fig. 1b). Additionally, the interaction between the treatments and parental genders also had a significant effect on parental care (χ² = 74.27, *P* < 0.001; Table 1). Specifically, compared to females and males in the contrasted group, males did not change their care in the resource availability group, whereas females increased their care (Fig. 1b). However, both males and females increased their level of care in the presence of intraspecific competition, but both decreased levels of care in the presence of interspecific competition (Fig. 1b).

**Figure 1.**
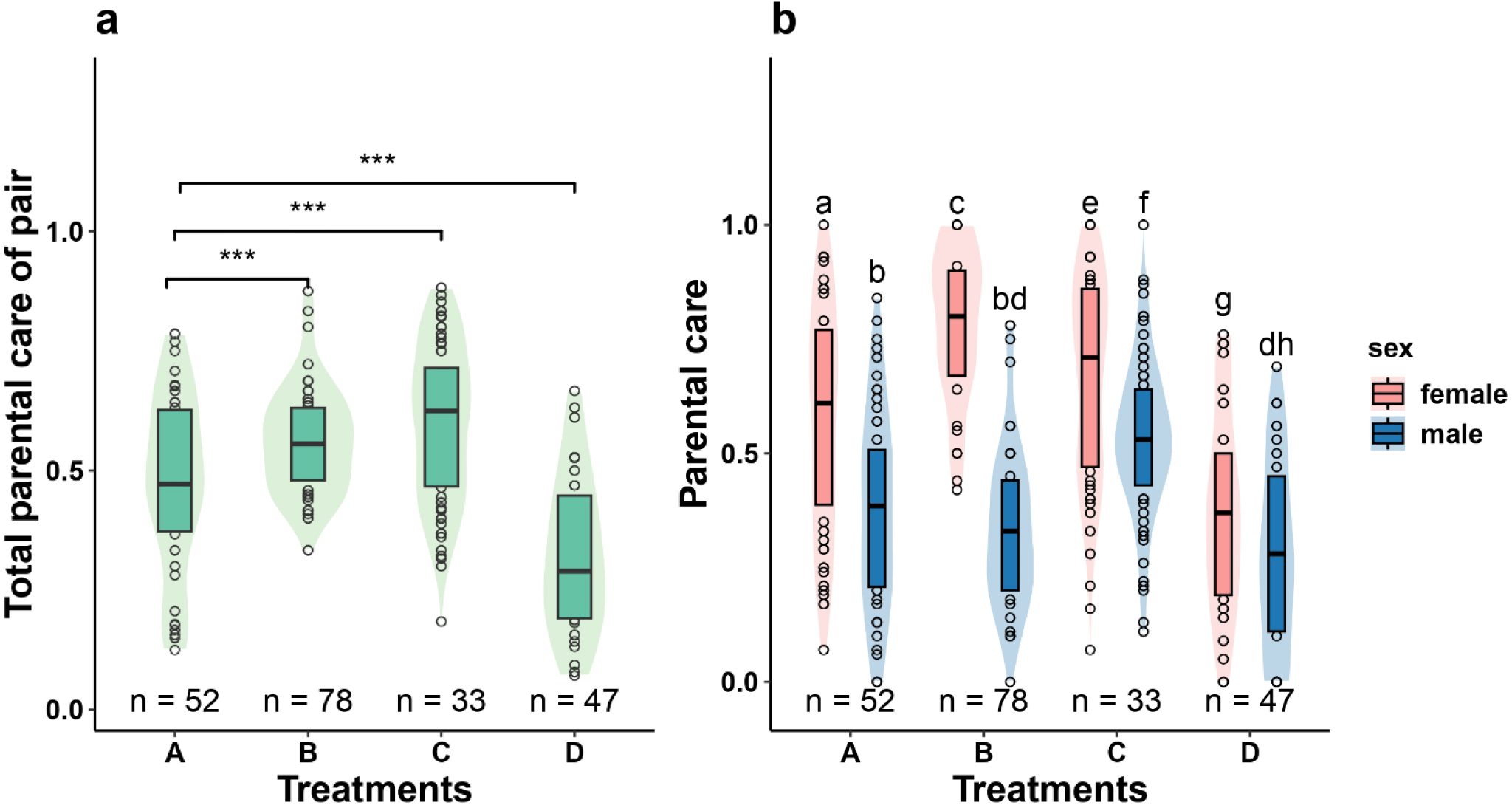
Comparison of (a) total parental care of pairs and (b) parental care between male and female burying beetles under experimental treatments (A: contrast, B: resource availability, C: intraspecific competition, D: interspecific competition). Blank circles indicate outliers, and ‘n’ represents sample sizes. The significance of the differences was determined using a *post hoc* pairwise test following the GLMs analysis. Significance code: ‘***’ is for P < 0.001. Identical letters in (b) indicate no significant difference in the means of parental care, while different letters indicate a significant difference.

**Table 1.** Difference of parental-care variables in the resource availability, intra- and interspecific competition groups compared with the contrasted group. Data in bold indicate statistically significant results (p < 0.05).

| Traits | Estimate | Standard error | Z-value | *P* value |
| --- | --- | --- | --- | --- |
| **Parental care of pair** |  |  |  |  |
| Resource availability | 0.32 | 0.09 | 3.76 | **< 0.001** |
| Intraspecific competition | 0.46 | 0.07 | 6.96 | **< 0.001** |
| Interspecific competition | -0.65 | 0.08 | -8.02 | **< 0.001** |
| **Parental care of individuals** |  |  |  |  |
| Resource availability | 1.04 | 0.19 | 5.45 | **< 0.001** |
| Intraspecific competition | 0.40 | 0.15 | 2.67 | **0.008** |
| Interspecific competition | -0.87 | 0.19 | -4.70 | **< 0.001** |
| Gender (baseline is female) | -0.83 | 0.11 | -7.53 | **< 0.001** |
| Resource availability × Gender | -1.20 | 0.19 | -6.23 | **< 0.001** |
| Intraspecific competition × Gender | 0.28 | 0.14 | 1.98 | **0.048** |
| Interspecific competition × Gender | 0.33 | 0.17 | 1.91 | 0.057 |
| **Ratio of male and female care** |  |  |  |  |
| Resource availability | -0.43 | 0.12 | -3.50 | **< 0.001** |
| Intraspecific competition | 0.19 | 0.09 | 2.03 | **0.042** |
| Interspecific competition | 0.08 | 0.13 | 0.65 | 0.519 |

The results of the GLM analysis for the ratio of male and female parental care showed a significance of experimental treatments (χ² = 33.05, *P* < 0.001). Compared to the contrasted group (mean ± SE = 0.85 ± 0.12), a significantly lower ratio of care between sexes was found in the resource availability group (mean ± SE = 0.47 ± 0.04; indicating a larger difference in parental care between sexes) while a significantly higher ratio was found in the intraspecific competition group (mean ± SE = 0.97 ± 0.10; indicating a weak difference on parental care between sexes) whereas no significant changes in the ratios were observed in the interspecific competition group (mean ± SE = 1.07 ± 0.30; Table 1, Fig.1b).

We found that both the treatments and spouse care had significant effects on the male and female care, whereas their interaction had not effect on male care but significant effect on female care (Table 2). In details, the GLM analysis for male care showed that 1) there were differences in male care between the contrasted goup and the three experimental groups (consistent with what we mentioned above); 2) female care had a significant positive effect on male care, which was independent from the treatments (Fig. 2a; although the slope of the male-female care trend line between the contrasted and the resource availability groups differs significantly, z = - 2.01, *P* = 0.044). Moreover, the GLM analysis for female care showed that 1) there were differences in female care between the contrasted goup and the three experimental groups (consistent with what we mentioned above); 2) the influence of male care on female care exhibited variability across experimental treatments. Females increased their care in response to increased male care in the contrasted and intraspecific competition groups, whereas their care remained unaffected by changes in male care in the resource availability and interspecific competition groups (Fig. 2b). There were significant differences in the slopes of the female-male care trend lines between the contrasted and the resource availability groups (z = - 2.19, *P* = 0.028), the contrasted and the intraspecific competiton groups (z = 2.15, *P* = 0.031).

**Figure 2.**
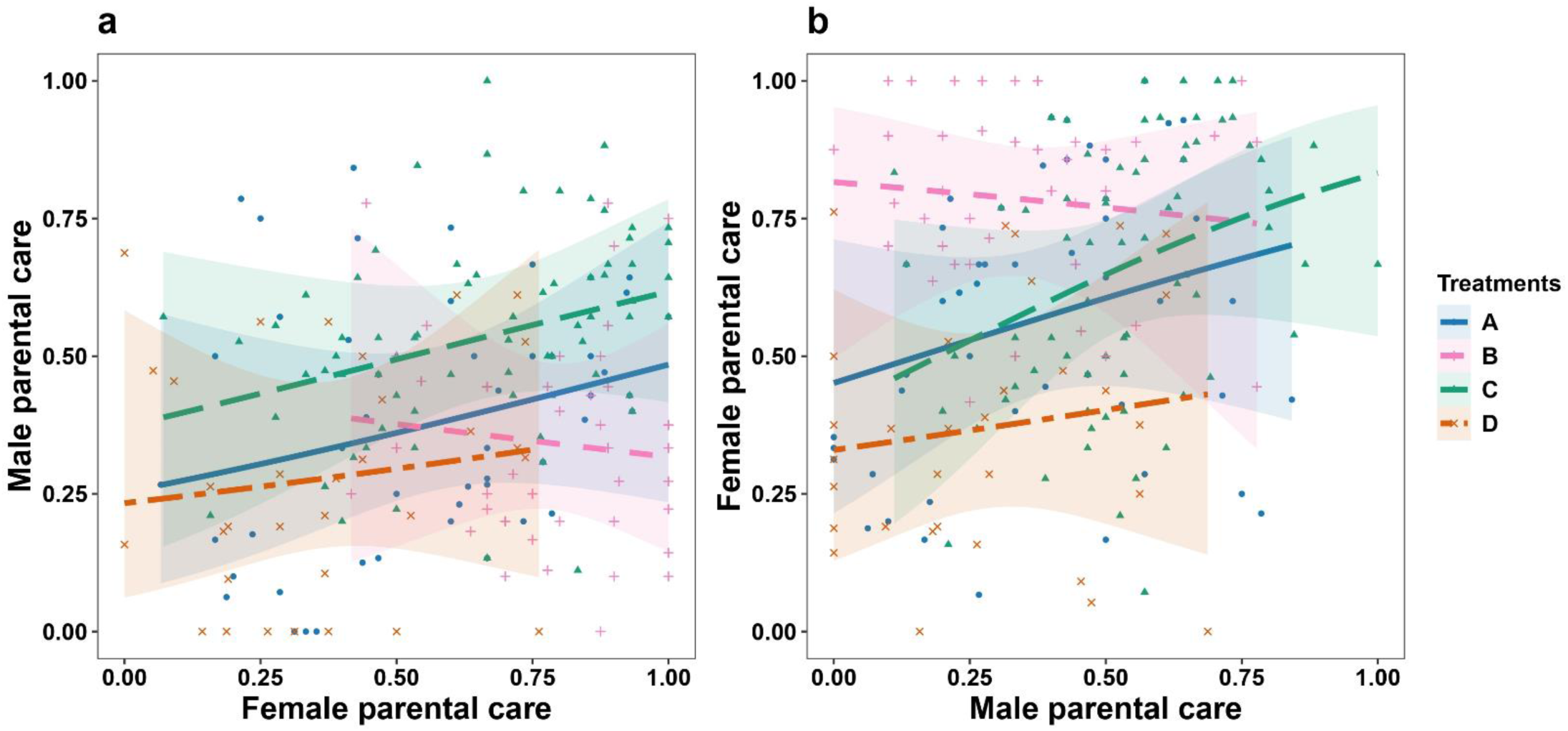
Males (a) and females (b) adjust their care respond to their spouse’s care in the treatments (A: contrast, B: resource availability, C: intraspecific competition, D: interspecific competition). The slops of trend lines obtained from the GLMs in (a) are as follows: A: Mean ± SE = 1.03 ± 0.34, z = 2.30, *P* = 0.01; B: Mean ± SE = -0.54 ± 0.63, z = -0.86, *P* = 0.86; C: Mean ± SE = 2.11 ± 0.36, z = 5.79, *P* < 0.001; D: Mean ± SE = 0.73 ± 0.42, z = 1.76, *P* = 0.28. In (b), the slopes are A: Mean ± SE = 0.89 ± 0.32, z = 2.77, *P* = 0.02; B: Mean ± SE = -0.49 ± 0.60, z = -0.81, *P* = 0.89; C: Mean ± SE = 1.06 ± 0.25, z = 4.20, *P* < 0.001; D: 0.75 ± 0.42, z = 1.78, *P* = 0.27.

**Table 2.** Effects of the experimental treatments (including contrasted), spouse care and their interaction on the male and female care. Data in bold indicate statistically significant results (p < 0.05).

| Explanatory variables | Male care |  |  | Female care |  |  |
| --- | --- | --- | --- | --- | --- | --- |
| | $\chi^2$ | df | *P* value | $\chi^2$ | df | *P* value |
| Treatments | 130.70 | 3 | **< 0.001** | 160.40 | 3 | **< 0.001** |
| Spouse care | 23.76 | 1 | **< 0.001** | 32.79 | 1 | **< 0.001** |
| Treatments × Spouse care | 5.69 | 3 | 0.128 | 15.21 | 3 | **0.002** |

### The influence of resource availability, intra- and interspecific competition on reproductive success and parental body mass change

Our results found that the experimental treatments had significant effects on brood size, brood mass and offspring number (Table 3). Compared to the contrasted groups, similar brood size, similar offspring number, but heavier dispersal larvae were found in the resource availability group (Fig.3 a, b and c). For the intraspecific competition group, all three components of reproductive success were similar as those for the contrasted group (Fig. 3 a, b and c). Conversely, the interspecific competition group exhibited smaller brood size, lighter brood mass, and fewer newly-eclosed beetles than that in the contrasted group (Fig. 3 a, b and c). We found that neither male nor female care influenced brood size and brood mass. However, female care, but not male care, had a significantly positive effect on offspring number (Table 3).

**Figure 3.**
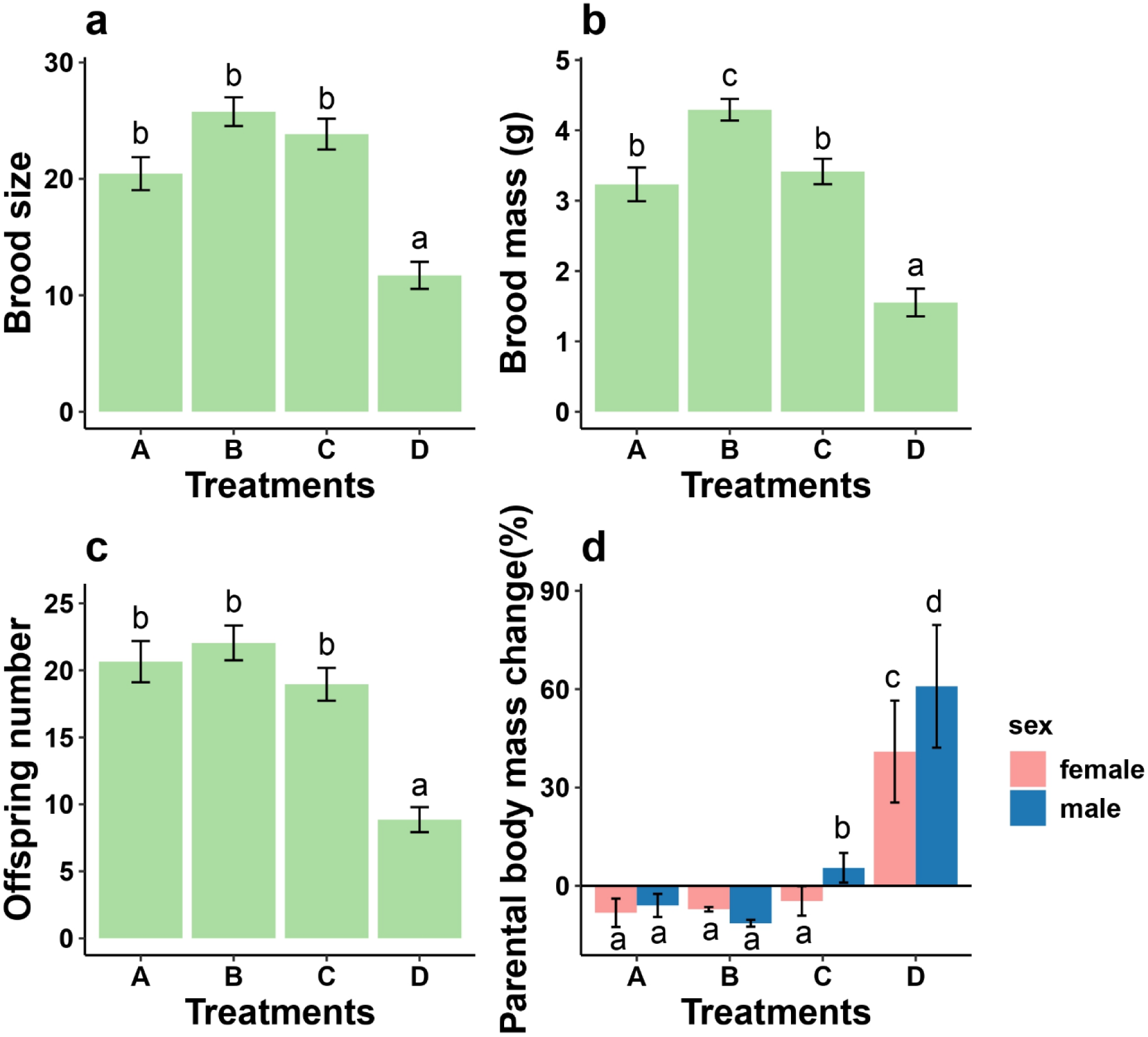
The difference in (a) brood size, (b) brood mass, (c) offspring number and (d) parental body mass change between the experimental treatments (A: contrast, B: resource availability, C: intraspecific competition, D: interspecific competition). Identical letters obtained from pairwise *post hoc* tests indicate no significant difference, while different letters indicate a significant difference.

**Table 3.** Effects of the experimental treatments (including contrasted) and parental care, on brood size, brood mass, and offspring number. Data in bold indicate statistically significant results (p < 0.05).

| Explanatory variables | Brood size |  |  | Brood mass |  |  | Offspring number |  |  |
| --- | --- | --- | --- | --- | --- | --- | --- | --- | --- |
| | $\chi^2$ | df | *P* value | *F* | df | *P* value | $\chi^2$ | df | *P* value |
| Treatments | 15.47 | 3 | **0.001** | 8.92 | 3 | **<0.001** | 13.08 | 3 | **0.004** |
| Male care | 1.02 | 1 | 0.312 | 2.71 | 1 | 0.102 | 0.10 | 1 | 0.753 |
| Female care | 2.78 | 1 | 0.095 | 2.10 | 1 | 0.15 | 4.30 | 1 | **0.038** |

We found from the LMM analysis that the experimental treatments (*F* = 18.87, *P* < 0.001) and parental genders (*F* = 10.81, *P* = 0.001) had significant effects on the parental body mass change, but the interaction between treatments and parental genders did not (*F* = 0.512, *P* = 0.674). Compared to individuals in contrasted groups, individuals in the interspecific competition group had more weight gain after breeding, whereas weight change was similar for individuals in the resource availability and the intraspecific competition groups (Fig. 3d). Furthermore, males obtained more weight after breeding than females (Fig. 3d). Specifically, individuals gained more weight with increasing level of their care (*F* = 6.09, *P* = 0.002).

## Discussion

Our study revealed significant effects of ecological and social environments in shaping variations in parental care and reproductive success among burying beetles. Specifically, compared to beetles that bred on large carcasses (the contrasted group), we found a higher level of total parental care in the resource availability group (breeding on small carcasses), which may be due to unchanged male care and increased female care. However, the presence of intraspecific competitors resulted in elevated parental care from both males and females. Conversely, both males and females reduced their parental care in the interspecific competition group (compared to the contrasted group), leading to lower total parental care. Our results also showed that males increased their care with the increasing level of female care, regardless of the ecological and social environments encountered during breeding. In contrast, females conditionally adjusted their care in response to increased male care, with heightened female care when breeding on large carcass and the intraspecific competition was present, while with unchanged female care when breeding on small carcasses and in the presence of interspecific competition. Subsequently, we found that our experimental manipulations significantly affected reproductive success, and parental body mass change during breeding. Beetle pairs breeding in the interspecific competition group produced the smallest brood size, lightest brood mass, and fewest offspring, but all parents obtained more body mass after breeding. In contrast, the other three groups did not differ significantly in these measures, except for the resource availability group, which exhibited a heavier brood mass compared to the control group. Female care had a significantly positive effect on the number of offspring. Additionally, both males and females that provided more care to offspring gained more body mass after breeding.

### Males and females adjusted parental care in relation to resource availability

Carcass availability (the extent of decomposition) and its size is the prior environmental pressure that determines burying beetles’ decisions on whether and how much they invest for their offspring, which also impacts the parental body maintenance and larvae growth directly. Large carcasses could support more breeders, and this also explain why communal breeding always occurs on larger carcasses in nature (Scott, 1998; Ma *et al*., 2022), because of no restricted access to the carcass for females and males due to the low level of sexual conflict. Meanwhile, larger carcasses could support more resources and energy for the development of larvae and their immunity (Smiseth *et al*., 2014; Shukla *et al*., 2018b). It has been known that female burying beetles can manipulate clutch size and brood size based on the size of carcass they occupy (Bartlett, 1987a; Müller *et al*., 1990b; Richardson & Smiseth, 2019). Based on this highly correlation between clutch size and brood size (we did not count the number off eggs), we would expect relatively more dispersal larvae (i.e., brood size), heavier brood mass and subsequently more newly-eclosed beetles (i.e., offspring number) for the pairs breeding on large carcasses (i.e., the contrasted group). However, this was not the case. Instead, we found a similar brood size, heavier brood mass, and similar offspring number for the pairs breeding on small carcasses (i.e., the resource availability group) compared to those on large carcasses, indicating more unhatched eggs or higher mortality of developing larvae before dispersal on large carcasses. The drawback of large carcasses is that they require breeding pairs to invest more time and energy in preparation, defence, and maintenance (Ma *et al*., 2022). Otherwise, poorly buried carcasses would harbour a high load of bacteria (e.g., fungi) that can transfer harmful pathogens to both adult beetles and developing larvae during their feeding activities on the carcass (Jacobs *et al*., 2014; Shukla *et al*., 2018a; Shukla *et al*., 2018b; Wang & Rozen, 2019; Salem & Kaltenpoth, 2022). The intensive microbial activities on large carcasses may partly explain the negative effects of large carcass on reproduction and offspring performance. In addition, consistent with our prediction, compared to the contrast group, females in the resource availability group increased their care for developing larvae, with heavier brood mass. Contrary to our prediction that males breeding on small carcasses would decrease their care, males in the resource availability group did not change their level of care compared to those in the contrasted group. One possible reason for the distinct responses of males and females to variations in carcass size is that females are relieved from the carcass maintenance duties when breeding on small carcasses. Although males and females have different responsibilities during breeding in this species, specifically in a biparental care system, females invest time and energy into offspring provisioning (direct care), whereas males focus on carcass maintenance and defending against intruders (indirect care) (Scott, 1990; Müller *et al*., 1998; Smiseth *et al*., 2005; Boncoraglio & Kilner, 2012) When breeding on large carcasses, males and females are required to maintain the carcasses by smearing the carcass with antimicrobial exudates, which contributes to delaying the decomposition of the carcasses (Duarte *et al*., 2018; Wang & Rozen, 2018). Conversely, when breeding on small carcasses, males can manage the workload of defending against intruders and maintaining the carcasses, allowing females to allocate more time and energy caring for their developing larvae.

In this species, female care rather than male care has greater effects on offspring fitness (e.g., Eggert *et al*., 1998; Müller *et al*., 1998; Smiseth *et al*., 2005), and females can also benefit from male desertion (Scott, 1990; Boncoraglio & Kilner, 2012). When breeding resources are limited, females may restrict males to access the carcass, whereas males may selectively abandon the carcass and larvae, ensuring these larvae to receive enough resources for development. However, we found that males consistently responded to the increase in female care by increasing their care in both the contrasted and resource availability groups, which supports the matching strategy hypothesis (Johnstone & Hinde, 2006). A positive relation between weight change of parents during breeding and their level of care suggests that access to the carcass benefits for parents because they could consume the carcass and save more energy for future reproduction (Wang *et al*., 2021).

Unlike females, this case is more common in males, which allows them to improve their body condition that invest in future mating opportunities (Ma *et al*., 2022). When resources are enough to support multiple individuals, the presence of males may not confer disadvantages on the fitness of offspring. However, when resources are limited, the presence of males intensifies competition over resources between parents and offspring. Therefore, we suggest that males may not choose to abandon carcasses initially but be coerced by females. More studies, such as the direct effect of females on offspring fitness by manipulating their access to the carcass, are still needed for testing the hypothesis. From the perspective of females, we found that they increased their care in response to the increase of male care in the contrasted group but provided their care independently with the change of male care in the resource availability group. This indicates that females may employ a matching strategy when breeding on large carcasses, but adopt a sealed bid strategy when breeding on small carcasses in response to the change of male care (Houston & Davies, 1985; Johnstone & Hinde, 2006). Different from male care, female care had a significantly positive effect on offspring number, which indicates that when breeding on large carcasses, females respond the change of male care by deploying the matching strategies, thereby improving the fitness of offspring. . The sealed bid strategy adopted by females when breeding on small carcasses may be explained by their tendency to operate near their physical limitations (Wang *et al*., 2021). Given that males and females adopted different strategies to respond to the regulation i. n the amount of care of their mates when breeding in small carcasses, the largest difference in parental care between sexes (i.e., the minimal ratio between male to female care) was observed in our resource availability group.

### The presence of intraspecific competition has positive effects on male and female parental care

As we mentioned above, communal breeding (i.e., co-breeding) is more likely to occur on a large carcass, on which more than two breeders with the same sex contribute to a joint brood(e.g., Bartlett, 1987a, Komdeur *et al*., 2013, Ma *et al*., 2022). Thus the past research on effects of conspecific competition on target parental care and reproductive success was introduce dead conspecific into breeding events (Ratz *et al*., 2022). This methods may weaken the actual the extent of intraspecific competition in nature. Thus in this study, we introduced smaller conspecifics 12h after the original beetle pair buried the carcass, simultaneously to prevent co-breeding and to simulate real pressure of intraspecific competition. The presence of intraspecific intruders may let the residents to invest more efforts for offspring, leading to an increased coordination in parental care between sexes (Müller *et al*., 1990a; Trumbo, 2006, Ratz *et al*., 2022). Our result shows that both males and females in the intraspecific competition group increased their care compared to the contrasted group. Based on our experimental setup, no individual was observed to escape from the breeding containers during breeding, which may increase the opportunity for the smaller pair access to the carcass for food and produce their own offspring. Moreover, it is possible that these two pairs of beetles copulate chaotically causing uncertain paternity and maternity in the developing larvae (Eggert & Müller, 1992; Müller *et al*., 2007; Ma *et al*., 2022). In addition, burying beetles cannot recognise their offspring but typically cannibalise these larvae before their own first larva hatches (Bartlett, 1987b; Müller & Eggert, 1990; Scott, 1998). Focal males and females likely spend more time accessing the carcass to guard their offspring and avoid the risk of infanticides committed by co-breeders. Hence, the focal males and females are expected to benefit from ensuring current reproductive success by increasing their care for developing larvae (Bartlett, 1987a; Benowitz *et al*., 2013; Ma *et al*., 2022). On the other hand, the presence of intraspecific individuals may indicate a high local population density and low future reproductive opportunities, with intense competition for limited breeding resources and high levels of extra-pair copulation (Bartlett, 1987a; Jenkins *et al*., 2000). Consequently, males may spend more time on carcasses for their own benefits, such as consuming the carcass and guarding their mates. Another possible explanation for the increasing care in both males and females is that good body condition (i.e., increased body weight) and/or having extended experience with offspring care after current breeding could help them outcompete others for future mating and breeding opportunities (e.g., Beeler *et al*., 2002; Wang *et al*., 2021). However, this speculation may need more evidence, as males and females would always be expected to increase their care whenever they have the opportunity to breed. We suggest further research to explore issues of reproductive allocation and mate choice in this species.

We found no difference in brood size, brood mass, and offspring number between the contrasted and the intraspecific competition group, suggesting that increased parental care may buffer the negative effects of the conspecific intruders on components of reproductive success. Otherwise, in the scenario of decreasing or even lacking care for developing larvae from the original pair that encountered alloparental adults, we would foresee high reproductive failure, as the latter pair would infanticide the larvae that are not their own offspring (Scott, 1998). Furthermore, both males and females adopted the matching strategy in response to the adjustment in care of their mates, which could generate not only the same fitness benefits for both sexes; but also improve their body condition and reproductive experience *per se*. The maximum ratio of male to female care was observed in the intraspecific competition group, indicating the smallest difference in care between males and females. We suggest that the occurrence of conspecific competitors during the breeding phase would enhance the cooperation in the reproductive investment of males and females but weaken the sexual conflict over parental care.

### The presence of interspecific competition has negative effects on male and female parental care

Blowflies are one of the most common competitors for burying beetles in nature (Scott, 1998; Tsai *et al*., 2020), because they always discover dead animals quicker than other species and then lay eggs on them, and these eggs hatch and develop quickly. Furthermore, these feeding activities of maggots consume parts of the carcass and may produce some secondary metabolites which make the carcass unpalatable for burying beetles (Vogel *et al*., 2017; Shukla *et al*., 2018a; Chemnitz *et al*., 2020). Therefore, when carcasses are usurped by maggots, beetle parents, especially males, must spend more time cleaning the maggots (i.e., killing the maggots directly or preserving the carcass with secretions), which decreases the time they can spend providing care for their offspring. Consistent with our prediction, males in the interspecific competition group decreased their care for offspring compared to those in the contrasted group. Contrary to our prediction, however, females in the interspecific competition group did not increase their care but instead decreased it compared to those in the contrasted group. Due to the parental division of labour in burying beetles, both males and females may decrease their own care, but for different purposes. On one hand, male may spend time and energy to ward off maggots, however, on the other hand, they may desert current broods earlier as maggots feeding behaviour relatively limits the availability of males feeding on the carcass (Pilakouta *et al*., 2016). From the perspective of females, the occurrence of blowfly maggots normally predicts a high reproductive failure, which allows them to decrease their parental care to current broods and then pursue future reproductive opportunities. The interspecific competition group had less reproductive benefits, such as smaller brood size and lighter brood mass, compared to the contrasted group, which may be explained by the decreased parental care. From another perspective, the presence of fewer larvae might in turn lead to the decrease in male and female care (Müller *et al*., 1990b; Sahm *et al*., 2023). It has been demonstrated that maggots always outcompete burying beetle larvae in the absence of their parents (Bartlett, 1987a; Chan *et al*., 2019), which may cause adverse effect to parental and offspring fitness in both a short and a long timescale (e.g., Bartlett, 1987a; Scott, 1998). In breeding groups where reproduction is successful, both males and females were suggested to allocate some their time and energy to dispel these maggots. In this case, they can achieve benefits from feeding on these maggots to increase their body mass and save for future reproductive opportunities. Our results on parental body mass change also support this, showing that both males and females in the interspecific competition group gained more body mass after breeding compared to those in the contrasted group. The introduction of fly maggots into the breeding containers caused dramatic reproductive failure in this study (i.e., only 7 out of 33 broods had larvae that survived to dispersal), which may lead to inaccuracies in the analysis of brood size, brood mass, and offspring numbers in the interspecific competition group. However, such dramatic reproductive failure precisely reflected the significantly negative effects of blowfly maggots on the reproductive success of burying beetles. Then males and females might benefit from abandoning their current broods earlier and providing less care for their larvae, by which they can save energy to pursue future reproductive opportunities.

Similar to the resource availability group, we found that males still adopted a matching strategy in response to the adjustment of female care, whereas females employed a sealed bid strategy in response to the adjustment of male care. This result demonstrates that the blowfly maggots consume lots of carcasses causing resource limitations for burying beetles. However, these resource constraints changed as the maggots grew, which might require males and females to adjust their behavioural strategies over time. Accordingly, we suggest that when the blowfly maggots were present, female burying beetles regulate their care based on the quality of the carcasses over time and the number of their developing larvae, whereas male burying beetles only adjust their care to match the care of their mates. As males always gain more body mass after breeding than females, the male matching strategy might be explained by that males tend to gain more benefits for themselves by consuming of the carcass, instead of investing more for offspring. This interpretation seems plausible because parental care in this study included both direct and indirect care, measured by the frequency of parental individuals being present on or in the prepared carcasses. More detailed behavioural observations are needed to further explore male investment strategies during breeding in this species.

## Conclusion

In conclusion, our results in *N. vespilloides* indicate that resource availability and both intra- and interspecific competition result in varied reproductive costs. Males and females accordingly adjusted their parental care in response to these ecological and social factors with sex-specific differences. Furthermore, males are more likely to a matching strategy in response to the variation of male care, whereas females adjust their investment strategies from matching to sealed bid responding to the male care change based on the variations of carcass size and the presence of interspecific competition respectively. Our findings suggest the roles of ecological and social environments in influencing individual reproductive behaviour and strategies. In this research, we did not control the number of larvae, which is for eliminating human disturbances during laboratory manipulations, but the brood size did impact the parental investments. We encourage future studies to manipulate brood size.

## Acknowledgments

This study was supported by a Ph.D. grant from the China Scholarship Council (CSC, 202106180024) to DM, an Ecology Fund of the Royal Netherlands Academy of Arts and Sciences (KNAWWF/807/19021) to LM, and NWO grants (854.11.003 and 823.01.014) to JK. And we thank Maaike A. Versteegh for her help during the data analyses.

## Author contributions

LM and JK contributed to the study conception and design. LM performed the experiments. DM analyzed the data and wrote the first draft of the manuscript. LM and JK led the editing of the manuscript. All authors read and approved the final manuscript.

## Disclosure

### Ethics

This study did not require ethical approval for animal care.

## References

Achorn, A.M. and Rosenthal, G.G. (2020) It’s Not about Him: Mismeasuring ‘Good Genes’ in Sexual Selection. Trends in Ecology & Evolution, 35(3), 206–219.

Arnqvist, G. and Rowe, L. (2005) Sexual Conflict. Princeton (NJ): Princeton University Press.

Bartlett, J. (1987a) The behavioural ecology of the burying beetle Nicrophorus vespilloides (Coleoptera: Silphidae). University of Edinburgh.

Bartlett, J. (1987b) Filial cannibalism in burying beetles. Behavioral Ecology and Sociobiology 21:179–183.

Bates, D., Maechler, M., Bolker, B. and Walker, S. (2015) Fitting Linear Mixed-Effects Models Using lme4. Journal of Statistical Software, 67(1), 1–48.

Beeler, A.E., Rauter, C.M. and Moore, A.J. (2002) Mate discrimination by females in the burying beetle *Nicrophorus orbicollis*: The influence of male size on attractiveness to females. Ecological Entomology, 27(1), 1–6.

Benowitz, K.M., Head, M.L., Williams, C.A., Moore, A.J. and Royle, N.J. (2013) Male age mediates reproductive investment and response to paternity assurance. Proceedings of the Royal Society B: Biological Sciences, 280:20131124.

Boncoraglio, G. and Kilner, R.M. (2012) Female Burying Beetles Benefit from Male Desertion: Sexual Conflict and Counter-Adaptation over Parental Investment. PLOS ONE, 7(2), e31713.

Bonduriansky, R., Maklakov, A., Zajitschek, F. and Brooks, R. (2008) Sexual Selection, Sexual Conflict and the Evolution of Ageing and Life Span. Functional Ecology, 22(3), 443–453.

Chan, S.-F., Shih, W.-K., Chang, A.-Y., Shen, S.-F. and Chen, I.-C. (2019) Contrasting forms of competition set elevational range limits of species. Ecology Letters, 22, 1668–1679.

Chemnitz, J., von Hoermann, C., Ayasse, M. and Steiger, S. (2020) The Impact of Environmental Factors on the Efficacy of Chemical Communication in the Burying Beetle (Coleoptera: Silphidae). Journal of Insect Science, 20(4), 3.

Clutton-Brock, T.H. (1991) The evolution of parental care. Princeton University Press.

De Gasperin, O., Duarte, A. and Kilner, R.M. (2015) Interspecific interactions explain variation in the duration of paternal care in the burying beetle. Animal Behaviour, 190, 97–107.

Duarte, A., Welch, M., Swannack, C., Wagner, J. and Kilner, R.M. (2018) Strategies for managing rival bacterial communities: Lessons from burying beetles. Journal of Animal Ecology, 87(2), 414–427.

Eggert, A.-K., Reinking, M. and Müller, J.K. (1998) Parental care improves offspring survival and growth in burying beetles. Animal Behaviour, 55(1), 97–107.

Eggert, A.-K. and Müller, J.K. (1992) Joint breeding in female burying beetles. Behavioral Ecology and Sociobiology, 31(4), 237–242.

Fox, J. and Weisberg, S. (2019) An R Companion to Applied Regression, Third edition. Sage, Thousand Oaks CA.

Gauthier, A.H. and Jong, P.W.de. (2021) Costly children: The motivations for parental investment in children in a low fertility context. Genus, 77(1).

Gonzalez-Voyer, A., Thomas, G.H., Liker, A., Krüger, O., Komdeur, J. and Székely, T. (2022) Sex roles in birds: Phylogenetic analyses of the influence of climate, life histories and social environment. Ecology Letters, 25(3), 647–660.

Grayson, P., Glassey, B. and Forbes, S. (2013). Does brood parasitism induce paternal care in a polygynous host? Ethology, 119, 489e495.

Harrison, F., Barta, Z., Cuthill, I. and Székely, T. (2009) How is sexual conflict over parental care resolved? A meta-analysis. Journal of Evolutionary Biology, 22(9), 1800–1812.

Hopwood, P.E., Moore, A.J., Tregenza, T. and Royle, N.J. (2016) Niche variation and the maintenance of variation in body size in a burying beetle. Ecological Entomology, 41(1), 96– 104.

Hothorn, T., Bretz, F. and Westfall, P. (2008) Simultaneous Inference in General Parametric Models. Biometrical Journal, 50(3), 346–363.

Houston, A.I., Székely, T. and McNamara, J.M. (2005) Conflict between parents over care. Trends in Ecology & Evolution, 20(1), 33–38.

Houston, A.I. and Davies, N.B. (1985) The evolution of cooperation and life history in the dunnock Prunella modularis. In: British Ecological Society, Vol. 25. Behavioural Ecology: Ecological Consequences of Adaptive Behaviour. (R.M. Sibly & R.H. Smith eds), pp. 471–487. Blackwell, Oxford.

Janicke, T., Häderer, I.K., Lajeunesse, M.J. and Anthes, N. (2016) Darwinian sex roles confirmed across the animal kingdom. Science Advances, 2, e1500983.

Jacobs, C.G.C., Wang, Y., Vogel, H., Vilcinskas, A., van der Zee, M. and Rozen, D.E. (2014) Egg survival is reduced by grave-soil microbes in the carrion beetle, *Nicrophorus vespilloides*. BMC Evolutionary Biology, 14(1), 208.

Jenkins, E.V., Morris, C. and Blackman, S. (2000) Delayed benefits of paternal care in the burying beetle *Nicrophorus vespilloides*. Animal Behaviour, 60(4), 443–451.

Johnstone, R.A. and Hinde, C.A. (2006) Negotiation over offspring care—How should parents respond to each other’s efforts? Behavioral Ecology, 17(5), 818–827.

Lee, V.E., Head, M.L., Carter, M.J. and Royle, N.J. (2014) Effects of age and experience on contest behavior in the burying beetle, *Nicrophorus vespilloides*. Behavioral Ecology, 25, 172– 179.

Lenth, R. (2023) emmeans: Estimated Marginal Means, aka Least-Squares Means. R package version 1.9.0.

Lessells, C.M. (2012) Sexual conflict. In The evolution of parental care. Oxford University Press.

Liker, A., Freckleton, R.P. and Székely, T. (2013) The evolution of sex roles in birds is related to adult sex ratio. Nature Communications, 4, 1587.

Long, X., Liu, Y., Liker, A., Weissing, F.J., Komdeur, J. and Székely, T. (2022) Does ecology and life history predict parental cooperation in birds? A comparative analysis. Behavioral Ecology and Sociobiology, 76, 92.

Komdeur, J., Schrama, M.J.J., Meijer, K., Moore, A J. and Beukeboom, L.W. (2013) Cobreeding in the Burying Beetle, *Nicrophorus vespilloides*: Tolerance Rather Than Cooperation. Ethology, 119(12), 1138–1148.

Ma, D., Lu, M., Cheng, Z., Du, X., Zou, X., Yao, X., et al. (2021) Male parent birds exert more effort to reproduce in two desert passerines. Avian Research, 12, 37.

Ma, L., Versteegh, M.A., Hammers, M. and Komdeur, J. (2022) Sex-specific influence of communal breeding experience on parenting performance and fitness in a burying beetle. Royal Society Open Science, 9(2), 211179.

McNamara, J.M., Gasson, C.E. and Houston, A.I. (1999) Incorporating rules for responding into evolutionary games. Nature, 401(6751), Article 6751.

Müller, J.K. and Eggert, A.-K. (1990) Time-dependent shifts between infanticidal and parental behavior in female burying beetles a mechanism of indirect mother-offspring recognition. Behavioral Ecology and Sociobiology, 27, 11–16.

Müller, J.K., Eggert, A.-K. and Dressel, J. (1990a) Intraspecific brood parasitism in the burying beetle, *Necrophorus vespilloides* (Coleoptera: Silphidae). Animal Behaviour, 40(3), 491–499.

Müller, J.K., Eggert, A.-K. and Furlkrȍger, E. (1990b) Clutch Size Regulation in the Burying Beetle *Necrophorus vespilloides* Herbst (Coleoptera: Silphidae). Journal of Insect Behavior, 3(2), 265–270.

Müller, J.K., Eggert, A.-K. and Sakaluk, S.K. (1998) Carcass maintenance and biparental brood care in burying beetles: Are males redundant? Ecological Entomology, 23(2), 195–200.

Müller, J. K., Braunisch, V., Hwang, W. and Eggert, A.-K. (2007) Alternative tactics and individual reproductive success in natural associations of the burying beetle, *Nicrophorus vespilloides*. Behavioral Ecology, 18(1), 196–203.

Paquet, M. and Smiseth, P.T. (2016) Maternal effects as a mechanism for manipulating male care and resolving sexual conflict over care. Behavioral Ecology, 27(3), 685–694.

Paquet, M., Wotherspoon, R. and Smiseth, P.T. (2017) Caring males do not respond to cues about losses in paternity in the burying beetle *Nicrophorus vespilloides*. Animal Behaviour, 127, 213– 218.

Parker, G.A. (1979) Sexual selection and sexual conflict. In Sexual selection and reproductive competition in insects (ed. M. S. Blum & N. A. Blum), pp. 123–166. London: Academic Press.

Pilakouta, N., Richardson, J. and Smiseth, P.T. (2015) State-dependent cooperation in burying beetles: parents adjust their contribution towards care based on both their own and their partner’s size. Journal of Evolutionary Biology. 28, 1965–1974.

Pilakouta, N., Richardson, J. and Smiseth, P.T. (2016) If you eat, I eat: Resolution of sexual conflict over consumption from a shared resource. Animal Behaviour, 111, 175–180

Pontzer, H. and McGrosky, A. (2022) Balancing growth, reproduction, maintenance, and activity in evolved energy economies. Current Biology, 32(12), R709–R719.

R Core Team (2022) R: A Language and Environment for Statistical Computing. R Foundation for Statistical Computing, Vienna.

Ratz, T., Stenson, S. and Smiseth, P.T. (2020) Offspring beg more toward larger females in a burying beetle. Behavioral Ecology, 31(5), 1250–1256.

Ratz, T., Leissle, L. and Smiseth, P.T. (2022) The presence of conspecific intruders alters the magnitude of sex differences in care in a burying beetle. Animal Behaviour, 194, 57–65.

Requena, G.S., Buzatto, B.A., Munguía-Steyer, R. and Machado, G. (2009). Efficiency of uniparental male and female care against egg predators in two closely related syntopic harvestmen. Animal Behaviour, 78, 1169e1176.

Richardson, J. and Smiseth, P.T. (2019) Effects of variation in resource acquisition during different stages of the life cycle on life-history traits and trade-offs in a burying beetle. Journal of Evolutionary Biology, 32(1), 19–30.

Royle, N.J., Smiseth, P. and Kölliker, M. (2012) The evolution of parental care: summary, conclusions and implications. In: M. Kölliker, M.J. Royle, P.T. Smiseth and N.J. Royle (Eds.) The evolution of parental care. Oxford, UK: Oxford University Press

Sahm, J., Conrad, T., Scheu, L. and Steiger, S. (2023) Brood size, food availability, and body size affects male care decisions and offspring performance. Ecology and Evolution, 13(6), e10183.

Salem, H. and Kaltenpoth, M. (2022) Beetle–Bacterial Symbioses: Endless Forms Most Functional. Annual Review of Entomology, 67(1), 201–219.

Sanz, J.J., Kranenbarg, S. and Tinbergen, J.M. (2000) Differential response by males and females to manipulation of partner contribution in the great tit (Parus major). Journal of Animal Ecology, 69:74–84.

Schwagmeyer, P.L., Mock, D.W. and Parker, G.A. (2002) Biparental care in house sparrows: negotiation or sealed bid? Behavioral Ecology, 13, 713–721

Scott, M.P. (1990) Brood Guarding and the Evolution of Male Parental Care in Burying Beetles. Behavioral Ecology and Sociobiology, 26(1), 31–39.

Scott, M.P. (1998) The ecology and behavior of burying beetles. Annual Review of Entomology, 43(1), 595–618.

Shukla, S.P., Plata, C., Reichelt, M., Steiger, S., Heckel, D.G., Kaltenpoth, M., et al. (2018a) Microbiome-assisted carrion preservation aids larval development in a burying beetle. Proceedings of the National Academy of Sciences USA, 115(44), 11274–11279.

Shukla, S.P., Vogel, H., Heckel, D.G., Vilcinskas, A. and Kaltenpoth, M. (2018b) Burying beetles regulate the microbiome of carcasses and use it to transmit a core microbiota to their offspring. Molecular Ecology, 27(8), 1980–1991.

Smiseth, P.T., Andrews, C.P., Mattey, S.N. and Mooney, R. (2014) Phenotypic variation in resource acquisition influences trade-off between number and mass of offspring in a burying beetle. Journal of Zoology, 293(2), 80–83.

Smiseth, P.T., Dawson, C., Varley, E. and Moore, A.J. (2005) How do caring parents respond to mate loss? Differential response by males and females. Animal Behaviour, 69(3), 551–559.

Smiseth, P.T. and Moore, A.J. (2008) Parental Distribution of Resources in Relation to Larval Hunger and Size Rank in the Burying Beetle *Nicrophorus vespilloides*. Ethology, 114(8), 789– 796.

Soulsbury, C.D. (2019) Income and capital breeding in males: Energetic and physiological limitations on male mating strategies. Journal of Experimental Biology, 222(1), jeb184895.

Steiger, S., Peschke, K., Francke, W. and Müller, J.K. (2007) The smell of parents: Breeding status influences cuticular hydrocarbon pattern in the burying beetle *Nicrophorus vespilloides*. Proceedings of the Royal Society B: Biological Sciences, 274(1622), 2211–2220.

Székely, T., Weissing, F.J. and Komdeur, J. (2014) Adult sex ratio variation: implications for breeding system evolution, Journal of Evolutionary Biology, 24(8), 1500–1512.

Taborsky, B., Guyer, L. and Demus, P. (2014) ‘Prudent habitat choice’: a novel mechanism of size-assortative mating. Journal of Evolutionary Biology, 27:1217–1228.

Trivers, R.L. (1972) Parental Investment and Sexual Selection. In Sexual Selection and the Descent of Man, ed. B. Campbell (Chicago: Aldine Press), 136–179.

Trumbo, S.T. (2006) Infanticide, sexual selection and task specialization in a biparental burying beetle. Animal Behaviour, 72(5), 1159–1167.

Trumbo, S.T. (2022) Why do males stay in biparental burying beetles? Behaviour, 159(13–14), 1301–1318.

Tsai, H.-Y., Rubenstein, D.R., Chen, B.-F., Liu, M., Chan, S.-F., Chen, D.-P., et al. (2020). Antagonistic effects of intraspecific cooperation and interspecific competition on thermal performance. eLife, 9, e57022.

Vági, B., Végvári, Z., Liker, A., Freckleton, R.P. and Székely, T. (2019) Parental care and the evolution of terrestriality in frogs. Proceedings of the Royal Society B: Biological Sciences, 286, 20182737.

van Dijk, R.E., Székely, T., Komdeur, J., Pogány, Á., Fawcett, T.W. and Weissing, F.J. (2012) Individual variation and the resolution of conflict over parental care in penduline tits. Proceedings of the Royal Society B: Biological Science. 279: 1927–1936.

Venables, W.N. and Ripley, B.D. (2002) Modern Applied Statistics with S. Fourth Edition. *Springer*, New York.

Vogel, H., Shukla, S.P., Engl, T., Weiss, B., Fischer, R., Steiger, S., et al. (2017) The digestive and defensive basis of carcass utilization by the burying beetle and its microbiota. Nature Communications, 8(1).

Wang, W., Ma, L., Versteegh, M.A., Wu, H. and Komdeur, J. (2021) Parental Care System and Brood Size Drive Sex Difference in Reproductive Allocation: An Experimental Study on Burying Beetles. Frontiers in Ecology and Evolution, 9.

Wang, W., Ma, L., Versteegh, M.A., Wu, H. and Komdeur, J. (2022) Detection of reproductive trade-offs is influenced by resource availability and maintenance: An experimental study in the burying beetle (*Nicrophorus vespilloides*). Behavioral Ecology and Sociobiology, 76(6), 76.

Wang, Y. and Rozen, D.E. (2018) Gut microbiota in the burying beetle, *Nicrophorus vespilloides*, provide colonization resistance against larval bacterial pathogens. Ecology and Evolution, 8(3), 1646–1654.

Wang, Y. and Rozen, D.E. (2019) Fitness costs of phoretic nematodes in the burying beetle, *Nicrophorus vespilloides*. Ecology and Evolution, 9(1), 26–35.

Wickham, H. (2016) ggplot2: Elegant Graphics for Data Analysis. Springer-Verlag, New York.

Zheng, J., Li, D. and Zhang, Z. (2018) Breeding biology and parental care strategy of the little-known Chinese Penduline Tit (*Remiz consobrinus*). Journal of Ornithology. 159, 657–666.

